# Celastrol directly inhibits PFKM to induce weight loss and leptin sensitization

**DOI:** 10.1101/2020.09.06.284752

**Authors:** Kang Wang, Xiaobo Wu, Yinghua Zhuang, Huan Sun, Fengchao Wang, Tao Wang, Zhiyuan Zhang

## Abstract

Despite the prevalence of obesity and related health consequences around the globe, effective treatments for inducing healthy weight loss are still lacking. Celastrol is a pentacyclic triterpene that was recently identified as a potent anti-obesity agent. Celastrol increases sensitivity to leptin, but the molecular target of celastrol is unknown. Therefore, the mechanisms by which this agent exerts its anti-obesity effect remain elusive. Using tissue-specific ABPP (activity-based protein profiling), we found that PFKM, a rate-limiting enzyme for glycolysis in skeletal muscle, is a direct target of celastrol. Celastrol inhibited PFKM enzymatic activity, and *Pfkm* knockout mice were resistant to a high fat diet, were hypersensitive to exogenous leptin, and were unresponsive to celastrol. PFKM inhibition led to activation of AMPK and inactivation of ACC in cultured myotubes and mouse skeletal muscle. Specific loss of AMPK in muscle significantly attenuated the anti-obesity effects of celastrol. Further, PFKM inhibition and subsequent activation of the AMPK/ACC signaling pathway reduced levels of free fatty acids by switching energy expenditure and consequently decreasing levels of SOCS1 expression, which are both required for leptin sensitization in 293t/hLepRb cells and mice. Finally, using a high throughput compound screen we identified an alternative PFKM inhibitor, 3-79, which exhibits a strong anti-obesity effect and non-covalent binding capacity. This compound is a promising agent for treating obesity in the clinic.

## Introduction

Obesity is a metabolic disorder involving abnormal or excessive fat accumulation, affecting more than 900 million people worldwide^1^. Obesity is becoming one of the major risk factors for a series of human diseases, including type 2 diabetes, cardiovascular disease, musculoskeletal disorders, and cancer. The adipocyte-derived hormone leptin functions as an anorexic hormone, regulating energy homeostasis, metabolism, and neuroendocrine function^2–4^. Levels of leptin therefore provide information about peripheral energy conditions. Leptin reduces body weight primarily by stimulating anorexigenic neuropeptides including POMC (proopiomelanocortin) and CART (cocaine- and amphetamine-regulated transcript), and by repressing orexigenic neuropeptides such as AgRP (agouti-related protein) and NPY (neuropeptide Y) in the CNS (central nervous system)^5–8^. Leptin can also directly increase catabolism of peripheral tissues^9–11^. Although leptin reduces food intake and body weight in lean mice, high levels of leptin fail to suppress food intake and stimulate energy expenditure in mouse models of DIO (diet-induced obesity) and in obese humans^2,12,13^. Because obese individuals develop resistance to leptin, leptin-based clinical therapies for treating DIO are not effective.

Through an *in silico* drug screen, Ozcan and his co-workers identified celastrol as a potential leptin sensitizer and anti-obesity agent. Testing in DIO mice revealed that celastrol suppresses food intake, increases energy expenditure, and reduces body weight^14^. It was later found that celastrol induces the expression of Interleukin 1 receptor 1 (IL1R1) in the hypothalamus, and that this mediates celastrol’s leptin sensitization and anti-obesity effects^15^. However, there is no evidence that celastrol directly binds IL1R1. Several putative targets of celastrol have been identified, including IkB kinase, Hsp90/CDC37, protein tyrosine phosphatase 1B (PTP1B), T-cell PTP (TCPTP), and the proteasome, but none of these mediate the anti-obesity effects^16–20^. Thus, the direct molecular targets by which celastrol regulates leptin sensitivity remain elusive.

Skeletal muscles of obese individuals exhibit dramatic increases in glycolytic rate and lipid content^21,22^. In addition, activity of PFK (6-phosphofructokinase), a rate-determining enzyme in glycolysis, in skeletal muscle is proportional to alterations in body weight^21,23^. There are three isoforms of PFK, namely the muscle type (PFKM), liver type (PFKL) and platelet type (PFKC). While most tissues express all three isoforms, skeletal muscles express only PFKM^24^.

Using ABPP (activity-based protein profiling), we identified PFKM as a direct target of celastrol. Celastrol inhibits PFKM, leading to activation of AMPK (Adenosine 5’-monophosphate-activated protein kinase), leptin sensitization, and anti-obesity effects. We further demonstrate that celastrol switches energy demand from glycolysis to free fatty acid oxidation in skeletal muscle, and that this metabolic change in skeletal muscle elevates both cellular and circulatory fatty acid levels. This systematically increases leptin sensitivity and protects animals from DIO.

## Results

To identify targets of celastrol that lead to leptin sensitivity, we utilized activity-based protein profiling (ABPP). We designed a celastrol-based probe, 1-85, and added a propargyl tag to the C_28_ of celastrol to click to a biotin tag for further enrichment (Figure 1A). We first tested the anti-obesity effects of 1-85 in mouse models of DIO using celastrol as a positive control. DIO mice treated with celastrol or 1-85 for 20 days exhibited reduced body weight compared to controls (Figures S1A and S1B). Upon co-treatment with leptin, both celastrol and 1-85 reduced body weight in 8 week-old mice fed with normal chew diet compared to treatment with leptin or celastrol/1-85 alone (Figure S1C). Importantly, these results demonstrated that the anti-obesity effects of celastrol are conserved in this probe, providing an opportunity to identify the desired target.

**Figure 1.**
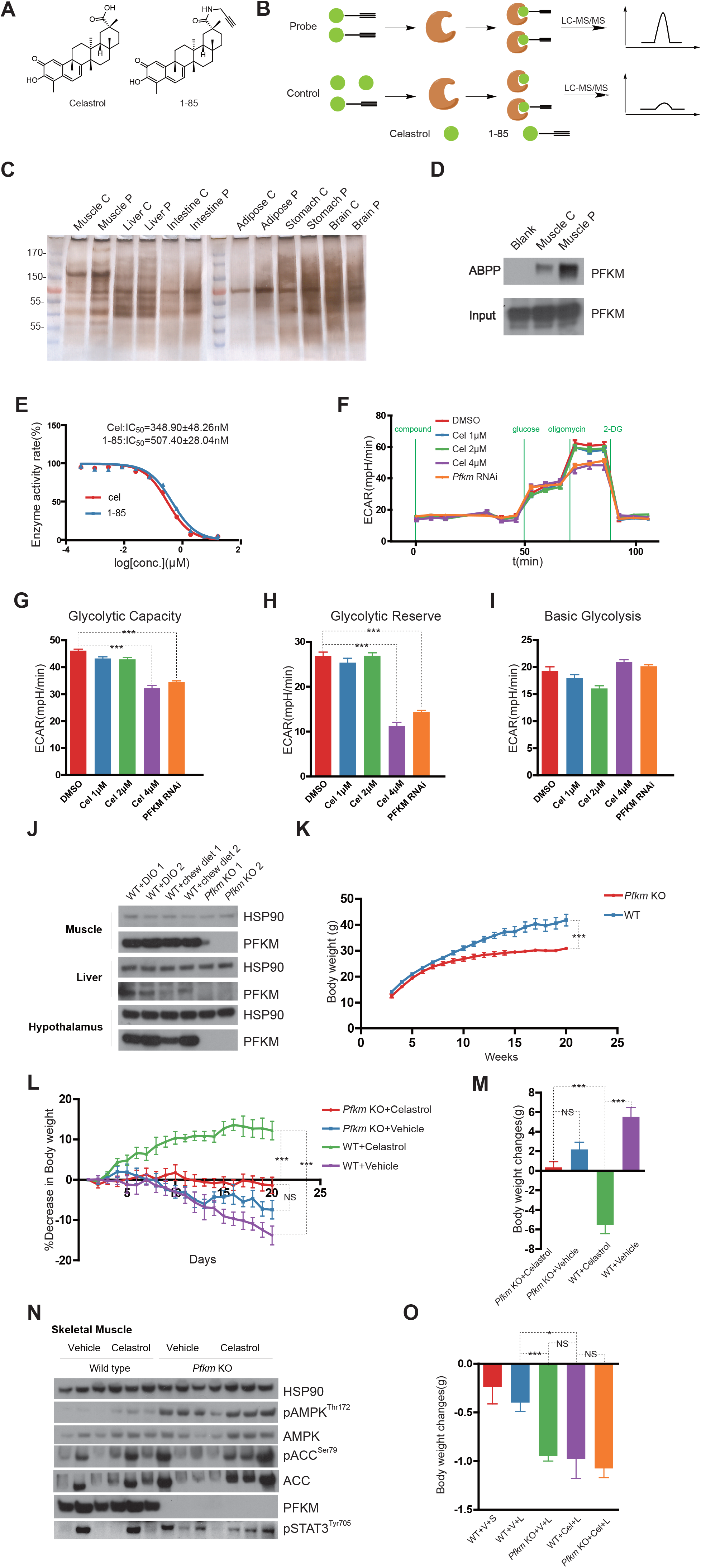
Celastrol causes weight loss by inhibiting PFKM. (A) Chemical structures of celastrol and 1-85. (B) Schematic diagram of ABPP. A 10-fold excess of celastrol was added to antagonize the 1-85 probe in controls. (C) The 1-85-bound proteins were analyzed by SDS-PAGE followed by silver staining. ABPP experiments were performed using homogenates of six organs: muscle, liver, intestine, adipose, stomach, and brain. (D) Western blot of 1-85-bound proteins in homogenates of skeletal muscle from 8-week-old lean mice using anti-PFKM antibodies. (E) The IC_50_ of celastrol and 1-85 was measured using recombinant PFKM. (F) Seahorse glycolysis stress test of L6 cells treated with celastrol or *PFKM* shRNA. (G-I) The glycolysis capacity (G), glycolytic reverse (H), and basic glycolytic rate (I) of each group in (F). (J) PFKM expression levels in muscle, liver, and hypothalamus from wild-type DIO mice, wild-type lean mice, and *Pfkm* KO mice. HSP90 was used as an internal control. (K) Body weight of wild-type and *Pfkm* KO mice fed a HFD. Mice were fed a HFD right after weaning for 20 weeks. Body weight of wild-type and *Pfkm* KO mice increased by 1.9-fold (from 14.16±2.47 g to 41.82±7.52 g; n=11) and 1.4-fold (from 12.55±1.94 g to 30.48±0.62 g; n=4), respectively. (L-M) Celastrol failed to reduce body weight of *Pfkm* KO DIO mice. Wild-type and *Pfkm* KO mice were fed HFD for 20 weeks to induce DIO, followed by daily intraperitoneal injection of celastrol (0.15 mg/kg) for 20 days. The time course of body-weight changes (L) and total body-weight changes (M) are shown. Celastrol reduced body weight of wild-type mice after 20-day treatment (from 45.8±2.10 g to 40.27±3.56 g, n=3), compared to vehicle treatment (from 40±1.57 g to 45.53±2.54 g, n=3). Neither vehicle (from 29.00±1.73 g to 31.2±2.98 g, n=3) nor celastrol (0.15 mg/kg) (from 29.15±2.36 g to 29.50±1.67 g, n=4) treatment reduced the body weight of *Pfkm* KO mice. (N) Western blot indicated that *Pfkm* KO and celastrol treatment induced phosphorylation of AMPK^Thr172^ and ACC^Ser79^ in skeletal muscles of DIO mice. (O) *Pfkm* KO mice were sensitive to leptin, but failed to further response to celastrol. Eight-week-old wild-type and *Pfkm* KO lean mice were intraperitoneally injected with either vehicle or celastrol (0.15 mg/kg) for four days. On the third and fourth days, mice were treated with either saline or leptin (5 mg/kg). Body-weight changes of WT control (−0.24±0.49 g; n=8), WT+L (−0.40±0.29 g; n=10), *Pfkm* KO+L (−0.95±0.10 g; n=4), WT+Cel+L (−0.975±0.40 g; n=4) and *Pfkm* KO+Cel+L (−1.08±0.19 g; n=4) between day 5 and day 3 were calculated.

Because the fundamental cause of obesity is imbalance between energy intake and expenditure, we performed ABPP using 1-85 as probe in six organs, namely muscle, liver, intestine, adipose, stomach, and brain (Figure 1B). Significant differences in protein bands between samples treated with probe vs. controls (samples treated with probe plus 10-fold of celastrol as a probe antagonist) were observed only in skeletal muscle (Figure 1C). Protein bands for both control and probe skeletal muscles were cut out and analyzed by LC-MS/MS, identifying a series of candidate proteins involved in metabolism (Table S1). Among these candidates was PFKM (Phosphofructokinase, Muscle), one of the three rate-determining enzymes in glycolysis. PFKM was one of the most enriched proteins and we confirmed this enrichment via western blot (Figure 1D). To determine direct effects of celastrol on enzymatic activity of PFKM, we ectopically expressed and purified the recombinant human PFKM in 293T cells, and found that both celastrol and 1-85 potently inhibited PFKM activity with IC50 values of 349 and 507 nM, respectively. This indicates that PFKM is a true target of celastrol (Figure 1E). We next performed seahorse glycolysis stress test using mature L6 myotube cells derived from skeletal muscle to examine if celastrol could reduce glycolysis by inhibiting PFKM *in vivo*. Both celastrol and *shRNA* (short-hairpin RNA) of *Pfkm* did not alter basal glycolytic rates measured by extracellular acidification rate (ECAR) (Figures 1F, 1I and S2), whereas adding oligomycin to block mitochondrial respiration rapidly increased the glycolytic rate (ECAR rate, 47.12±3.56 mpH/min) in control cells. Treatment with celastrol (ECAR rate, 30.58±6.43 mpH/min) and *Pfkm* shRNA (ECAR rate, 32.96±3.10 mpH/min) reduced this increase in glycolysis (Figures 1F and 1H). These data demonstrated that inhibition of PFKM by celastrol in L6 cell reduced the overall glycolytic capacity without affecting the basal glycolytic rate, suggesting that celastrol may only inhibit glycolysis in skeletal muscle when glycolytic rates are high. This could explained the previous results that celastrol reduced body weight only in DIO or leptin administered mice, since it has been demonstrated that obesity and leptin treatment increases glycolysis rate in skeletal muscle^3,21,23^.

To verify that PFKM is a *bona fide* target of celastrol in mediating weight loss, we generated *Pfkm* knockout (*Pfkm* KO) mice. Considering that PFKM deficiency might cause embryonic lethality, we attempted to generate Pfkm conditional knockout mice using Cre-loxP system (*Pfkm* KO, See method and Figure S3). However, homozygous *Pfkm* KO mice totally lacked PFKM expression in skeletal muscle, liver, and hypothalamus without Cre recombinase expression, which might because that the loxP insertions disrupt the transcription of *Pfkm* (See method and Figure 1J). Although *Pfkm* KO mice showed no obvious differences in behavior or development compared to wild-type littermates, *Pfkm* KO mice gained weight much more slowly than controls when fed a HFD (high fat diet). In addition, serum leptin levels of *Pfkm* KO mice were lower than their wild-type littermates (Figures 1K and S4A). After 20 weeks of HFD feeding, both *Pfkm* KO and wild-type mice were subjected to intraperitoneal administration of celastrol for 20 days. While wild-type mice lost body weight upon treatment with celastrol, body weight of *Pfkm* KO mice was unaffected, indicating that *Pfkm* mutant mice failed to response to celastrol (Figures 1L and 1M). To investigate if PFKM inhibition mediates the effects of celastrol on leptin sensitization, we treated both 8-week-old wild-type and *Pfkm* KO mice with leptin with or without celastrol. While wild-type mice treated with leptin showed mild effects in body weight, *Pfkm* KO mice showed significant reductions in body weight compared to wild-type controls. However, wild-type mice exhibited dramatic reductions in body weight upon co-injection of celastrol and leptin. By contrast, celastrol did not further reduce the body weight of leptin treated *Pfkm* KO mice (Figure 1O). These data indicated that *Pfkm* KO mice are more sensitive to leptin, and support that PFKM is a direct target of celastrol in promoting weight loss.

Considering that PFKM is a key regulator of glycolysis, pharmacological inhibition may reduce the contribution of energy from glycolysis. This would force cells to utilize other energy-generating pathways and thus exert profound influences on metabolic networks^25^. We then performed an unbiased, untargeted metabolomics analysis of an L6 cell line treated with celastrol (Table S2). Although ATP levels were unchanged, both ADP and AMP levels increased significantly after celastrol treatment (Figure 2A). In addition, malonyl-CoA levels were decreased (Figure 2A). Elevated levels of ADP and AMP could potentially activate AMPK (Adenosine 5’-monophosphate-activated protein kinase), which inhibits ACC (Acetyl-CoA carboxylase) catalyzing carboxylation of acetyl-CoA to produce malonyl-CoA. Therefore, we checked the activated states of AMPK, and found that celastrol treatment activated AMPK (phosphorylation of AMPK on thr172 is an activated form) and inactivated ACC (phosphorylation of ACC on ser790 is an inactivated form) in the L6 cell line (Figure 2B). AICAR (an AMPK activator) or acute glucose starvation had similar effects on the AMPK/ACC pathway. Consistent with a reduction in malonyl-CoA, which is a key intermediate in fatty acid synthesis and inhibits the beta-oxidation of fatty acids, the concentration of FFA (free fatty acids) was lower in L6 cells treated with celastrol (Figure 2C and table S3). We next verified that celastrol activated AMPK *in vivo*. Celastrol treatment of wild-type DIO mice significantly increased phosphorylation forms of AMPK^Thr172^ and ACC^Ser79^ in skeletal muscle but not in liver or hypothalamus (Figure 2F, 2G and 2H). Moreover, phosphorylation of AMPK was increased in muscle of *Pfkm* KO mice raised on HFD; this was not further increased by celastrol (Figures 1N, S5A and S5B). To investigate if the AMPK/ACC pathway in skeletal muscle mediates the effects of celastrol on weight loss, we specifically knocked out the α subunit of AMPK in skeletal muscle of *AMPKα1α2*^*loxp/loxp*^;*CKM-CreER* (*AMPKα1α2* mKO) mice. When fed a HFD for 20 weeks, *AMPKα1α2* mKO mice had same weight as controls. Leptin level in DIO *AMPK α1α2* mKO mice were equivalent to wild-type (Figure S4B). Importantly, loss of AMPK in muscles significantly attenuated the weight-loss effects of celastrol (Figures 2D and 2E). These results suggest that activation of AMPK is required for the weight-loss effects of celastrol.

**Figure 2.**
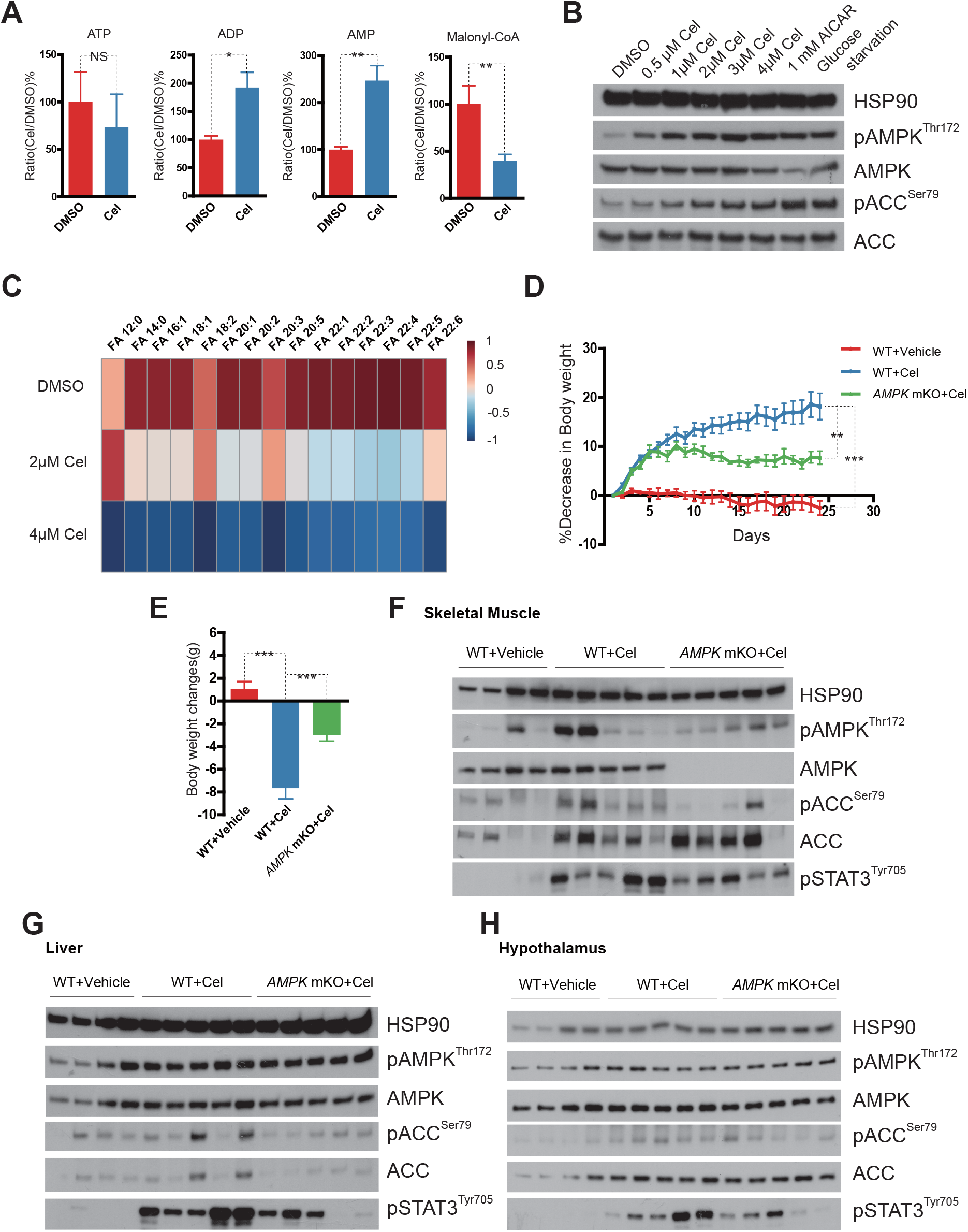
Celastrol reduces body weight and elevates systemic leptin response by activating AMPK in skeletal muscles. (A) The changes in ATP (from 100±55.51% to 73.19±60.46%), ADP (from 100±11.74% to 192.65±46.53%), AMP (from 100±10.63% to 247.09±55.43%), and Malonyl-CoA (from 100±38.49% to 39.54±14.15%) levels in L6 cell after 4 μM celastrol treatment for 30 min (n=3). (B) Western blot of L6 cells treated with different concentrations of celastrol for 30 min or 1 mM AICAR or glucose starvation for 8 h. (C) Heatmap visualization of relative free fatty acid levels in L6 cells after treatment with 2 μM or 4 μM celastrol for 30 min. The detailed data listed in table S3. (n=3 for each group) (D-E) Time course of percentage of body-weight changes (%) during compound treatment (D) and body weight changes (g) after 25 days of treatment (E) of wild-type or *AMPK*α*1*α*2* mKO DIO mice (n=4 for vehicle treated wild-type mice, n=5 for 0.15 mg/kg celastrol treated wild-type or *AMPK*α*1*α*2* mKO mice). (F-H) Western blot shows that celastrol induced phosphorylation of STAT3^Tyr705^ in skeletal muscles (F), liver (G), and hypothalamus (H), and *AMPK*α*1*α*2* mKO attenuated the effects of celastrol. Both wild-type and *AMPK*α*1*α*2* mKO DIO mice were intraperitoneally injected with celastrol (0.15 mg/kg) for 25 days.

We next examined whether celastrol-induced leptin sensitization is also mediated by the activation of AMPK. STAT3 activation (phosphorylation of STAT3 on Tyr705) is one of the most important downstream targets of leptin^26^, and we observed robust phosphorylation of STAT3 after acute intraperitoneal leptin administration in both the hypothalamus and skeletal muscle (Figure S6). We therefore used levels of STAT3^Tyr705^ phosphorylation as an indicator of leptin sensitivity in all tissues we investigated. Celastrol treatment increased STAT3^Tyr705^ phosphorylation in all tissues examined including liver, hypothalamus, and skeletal muscles of wild-type DIO mice, despite the fact that activation of AMPK was only detected in muscles. In contrast, levels of STAT3^Tyr705^ phosphorylation were higher in muscles of *Pfkm* KO mice. This was not further increased by celastrol (Figure 1N). Moreover, muscle-specific knockout of *AMPKα1α2* attenuated the ability of celastrol to induced STAT3^Tyr705^ phosphorylation in all these tissues (Figures 2F-2H).

Since inhibiting PFKM in muscles via celastrol increased cellular responses to leptin in most tissue, changes in circulating factors may mediate this systemic leptin sensitization. Previous research has demonstrated that obesity is associated with elevated levels of circulating fatty acids^27^. Based on the dramatic reductions in FFA in L6 cell after celastrol treatment, we assumed that high levels of FFA may interfere with the leptin signaling pathway. We then used a previously reported leptin-sensitive 293T cell model (293T/hLepRb) in which the human signaling form of the leptin receptor (hLepRb) is stably transfected^28^. We observed that extremely high levels of celastrol could inhibit both PFKM and PFKL (IC_50_=916.40 ± 35.66 nM), the dominant isoform of phosphofructokinase in 293T cell (Figure S7A). Celastrol treatment inhibited glycolytic capacity and reserve of 293T/hLepRb cells in seahorse glycolysis stress assay (Figure S7B). Treatment of 293T/hLepRb cells with 10 ng/ml leptin was sufficient to induce phosphorylation of STAT3^Tyr705^, and pretreating cells with palmitic acid (PA) attenuated leptin-induced STAT3 phosphorylation in a dose dependent manner (Figure 3A). Importantly, treatment with celastrol or AICAR, as well as glucose starvation, dramatically counteracted the suppressing effects of PA on leptin-induced phosphorylation of STAT3. Moreover, etomoxir, which reduces fatty acid oxidation by inhibiting carnitine palmitoyltransferase-1 (CPT-1), abolished the effects of celastrol, AICAR, or glucose starvation on leptin signaling (Figure 3B and 3C). These results, together with the fact that celastrol promotes consumption of FFA, indicate that celastrol sensitizes leptin signaling by reducing cellular FFA.

**Figure 3.**
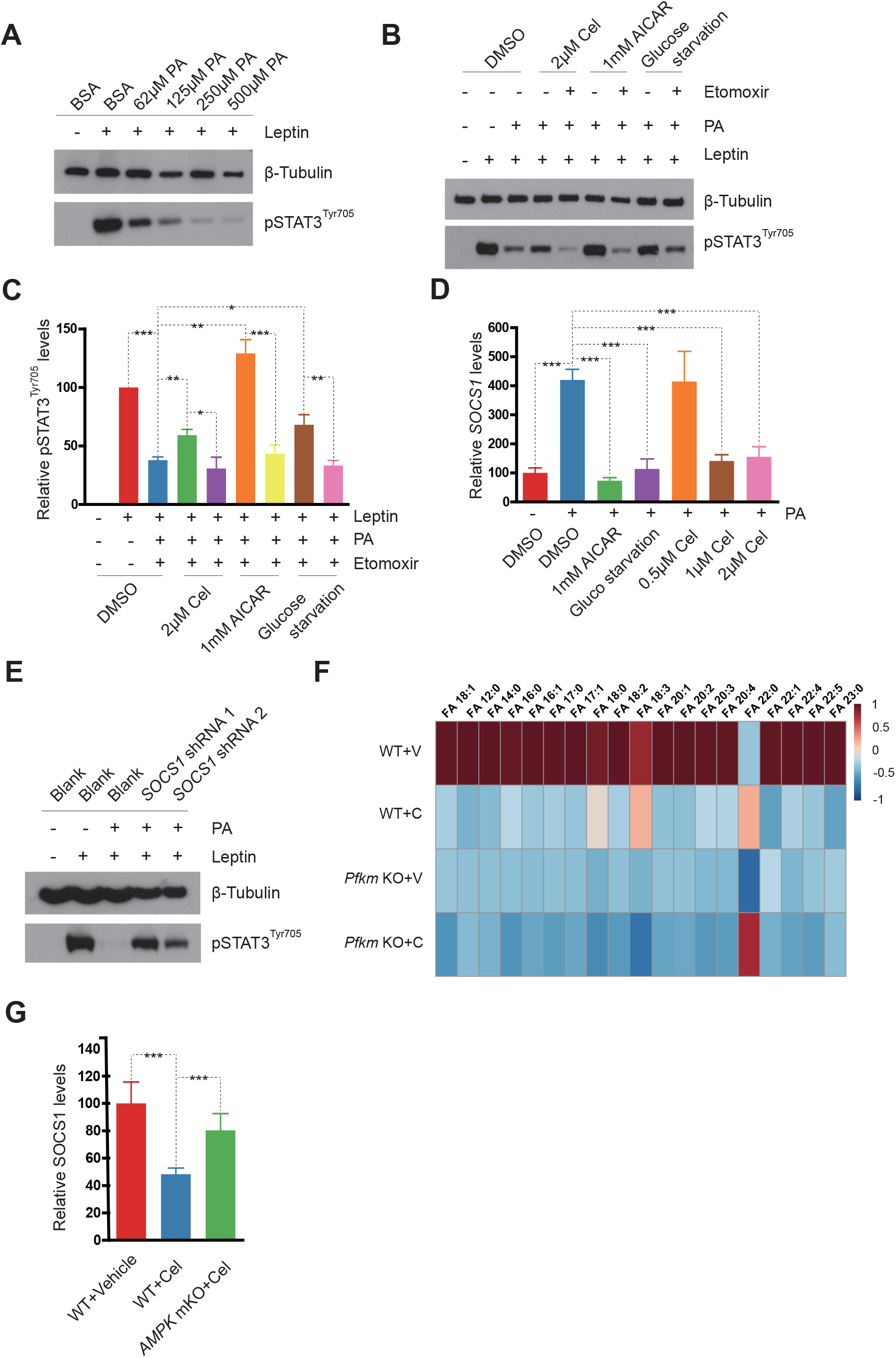
Reduction of FFA by celastrol is involved in leptin sensitization. (A) Western blot analysis shows PA attenuated the response of 293T/hLepRb cells to leptin. The 293T/hLepRb cells were pretreated with different concentration of PA for 8 h, followed by treated with 10 ng/ml leptin for 1 h. (B-C) Celastrol suppressed the inhibitory effects of PA on leptin response. The 293T/hLepRb cells were pretreated with 2 μM celastrol or 10 mM AICAR, or deprived of glucose for 8 h, followed by treatment with 500 μM PA for 8 h and subsequent treatment with 10 ng/ml leptin for 1 h. (C) Quantification of relative pSTAT3^Tyr705^ levels. β-Tubulin served as internal control (n=3). (D) QPCR was used to measure relative SOCS1 expressing levels in 293T/hLepRb cells co-treated with celastrol or AICAR or glucose starvation and 500 μM PA (n=3). (E) Western blot shows that 293T/hLepRb cells with SOCS1 knocked-down maintained leptin response in the presence of PA. The 293T/hLepRb cells were transfected with *SOCS1* shRNA expressing vector for 36 h, followed by treatment with 500 μM PA for 8 h and subsequent treatment with 10 ng/ml leptin for 1 h. (F) Lipidomic analysis of FFA changes in serum of WT and *Pfkm* KO mice intraperitoneal treated with celastrol or vehicle for 20 days (n=4 for *Pfkm* KO mice+celastrol, n=3 for the other groups). The numerical data is listed in Table S4. (F) SOCS1 expression levels in muscles of wild-type or *AMPK*α*1*α*2* mKO DIO mice. Both the wild-type and *AMPK*α*1*α*2* mKO mice were fed HFD for 20 weeks to induce DIO, followed by intraperitoneal injection of celastrol (0.15 mg/kg) for 20 days.

The SOCS family of proteins play important roles in the negative regulation of leptin signaling. We therefore reasoned that PA treatment could increase expression of SOCS1 and/or SOCS3. Since SOCS3 mRNA was not detected in 293T/hLepRb cells (data not show), we only measured the expression of SOCS1. SOCS1 was expressed more than 4-fold higher after 293T/hLepRb cells were treated with PA, and co-treatment with AICAR or glucose starvation reduced the expression of SOCS1 to levels without PA. Similarly, celastrol counteract the ability of PA to reduce expression of SOCS1 in a dose dependent manner (Figure 3D), suggesting that SOCS1 is a key regulator of PA-induced leptin desensitization. We next knocked down SOCS1 in 293T/hLepRb cells by *shRNA*, and found that these cells maintained phosphorylation of STAT3 upon PA treatment (Figures 3E and S8). We next verified these finding in DIO mice models, and found that total levels of circulating FFAs and most individual FFA species were largely decreased in wild-type mice following treatment of celastrol or *Pfkm* KO mice. By contrast, administration of celastrol to *Pfkm* KO mice showed no further reduction effects (Figure 3F and Table S4). Celastrol treatment significantly reduced SOCS1 mRNA levels in muscles, while muscle-specific *AMPK* knock-out attenuated the effect of celastrol on SOCS1 mRNA levels (Figure 3G). These results suggest that a reduction in circulatory FFAs mediates the effects of celastrol on leptin sensitization.

The anti-obesity effects of celastrol (by inhibiting PFKM in muscles), together with previous studies demonstrating that celastrol has excellent pharmacokinetic properties, indicate that celastrol may be a potent anti-obesity drug^29^. However, celastrol covalently binds to multiple targets leading to adverse side effects, thereby weakening the potential usage of celastrol as an anti-obesity drug^30,31^. Because celastrol is a complicated natural product, its pharmaceutical properties are difficult to improve. We therefore screened for alternative PFKM inhibitors from ~6000 pilot compounds (methods and supporting info). The twelve most potential compounds were selected for further analysis (Table S5). We measured the IC_50_ of these twelve compounds to PFKM in buffers with either 0.5 mg/ml bull serum albumin or 1 mM glutathione to exclude nonspecific protein binding and oxidizing, respectively. One of compounds, 3-79, inhibited PFKM in both systems (Figures 4A, 4B and Table S5). Seahorse glycolysis stress test revealed that 1 μM 3-79 reduced glycolytic capacity and reserve with minimal effects on the basal glycolytic rate (Figures 4C-4F) in L6 cells (as seen with celastrol). Moreover, 3-79 increased phosphorylation of AMPK^Thr172^ and ACC^Ser79^ in both L6 cells and skeletal muscles of DIO mice, and boosted leptin response in muscles (Figures 4G and 4H). Since 3-79 is an alternative inhibitor of PFKM, we next asked if 3-79 could induce anti-obesity effects in animal models. Intraperitoneal injection of 3-79 gradually reduced the body weight of DIO mice at comparable level as celastrol (Figures 4I and 4J). As with celastrol, injection of 3-79 at higher concentration did not have obvious effects on regular lean mice, indicating minimal toxicity effects of 3-79 (Figure S9). Considering that 3-79 has no covalent binding capacity and modification potential, it represents a potential PFKM inhibitor that should be further optimized.

**Figure 4.**
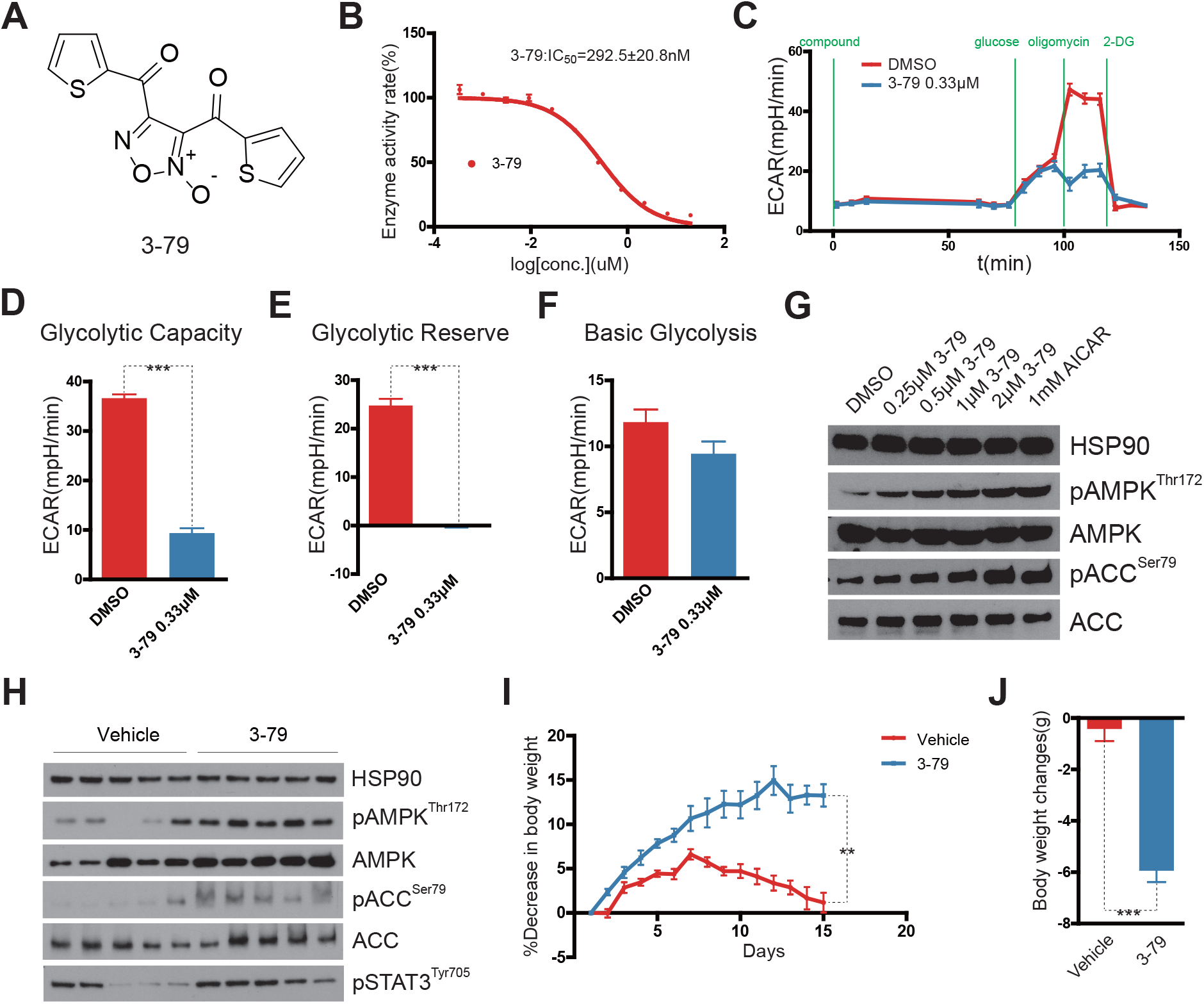
Identification of 3-79, a new PFKM inhibitor, as a potent candidate for pharmaceutic application. (A) Chemical structural of 3-79. (B) The IC_50_ of 3-79 on recombinant PFKM. (C-F) Seahorse glycolysis stress test shows that both glycolytic capacity (D, DMSO, 36.65±3.00 mpH/min n=5; 3-79, 9.40±4.11 mpH/min, n=6) and glycolytic reserve (E, DMSO, 24.80±5.22 mpH/min n=5; 3-79, −0.04±1.52 mpH/min, n=6) were reduced by 3-79 treatment. The ECAR of basal glycolysis (F, DMSO, 11.85olysismpH/min n=5; 3-79, 9.44±3.96 mpH/min, n=6) remained unchanged. (G-H) Western blot analysis of L6 cells (G) and skeletal muscle of DIO mice (H) reveal that 3-79 increased phosphorylation of AMPK^Thr172^, ACC^Ser79^ and STAT3^Tyr705^. L6 cells were treated with different concentrations of 3-79 or 1 mM AICAR for 30 min. The wild-type DIO mice were intraperitoneally injected with 3-79 (0.5 mg/kg) twice a day for 15 days. (I-J) 3-79 reduced body weight of wild-type DIO mice. Wild-type mice fed HFD for 20 weeks to induce DIO, followed by twice a day intraperitoneal injection of vehicle or 3-79 (0.5 mg/kg) twice a day for 15 days. The time course of relative body-weight changes (I, n=5) and body-weight changes after a 15-day period of 3-79 treatment (J, vehicle, Day1 41.63±1.51 g vs. Day 15 41.18±2.17 g; 3-79, Day1 41.66±1.79 g vs. Day15 36.82±2.26 g) are shown.

## Discussion

The discovery that leptin regulates food intake and energy expenditure heightened our understanding of bodyweight regulation. However, the attempt to cure obesity with leptin failed in most obese individuals that were not genetically deficient in leptin signaling. Circulating levels of leptin were extremely high in these obese patients, and the addition of exogenous leptin failed to decrease appetite and increase energy expenditure^12,13^. The cause of this leptin resistance has not been fully illuminated, but inflammation^32,33^, ER (endoplasmic reticulum) stress^34^, and autophagy^35^ are involved in obesity-induced leptin resistance. The discovery that celastrol can induce leptin sensitization, thereby reducing food intake and body weight in DIO mice with little effect on lean animals, re-established hope that leptin-oriented strategies could be used to treat obesity^14^. Celastrol treatment led to a significant decrease in PERK phosphorylation, but it fails to increase ER stress genes involved in hypothalamic ER stress, indicating that celastrol may not be directly involved in ER stress signaling^20^. Using a celastrol-based ABPP probe, named 1-85, we revealed that PFKM is a target of celastrol in skeletal muscle. Celastrol inhibited PFKM activity *in vitro*, and thus reduced glycolytic capacity in skeletal muscle-derived L6 myotube cells. *Pfkm* KO mice did not gain weight when fed a HFD, and did not lose weight in response to celastrol treatment. Moreover, the alternative PFKM inhibitor, 3-79, exhibited anti-obesity effects that mimicked celastrol. Altogether, PFKM is a major target of celastrol in promoting weight loss.

Leptin is a key regulator of fatty acid metabolism in skeletal muscle tissues, as this hormone stimulates fatty acid oxidation in muscle, primarily by activating the AMPK/ACC axis^36,37^. High level of glycolysis have been observed in both obese individuals and animals with leptin treatment^3,21^. As a rate-limiting enzyme in glycolysis, the activity of PFK is positively correlated with body weight gain^23^. As an inhibitor of glycolysis, celastrol increased cellular AMP levels, activating AMPK and inactivating ACC by phosphorylating ACC^Ser79^ in L6 cells, thereby promoting fatty acid oxidation. Therefore, celastrol-treated skeletal muscle cells produced more ATP from fatty acid oxidation rather than glycolysis. Similarly, *Pfkm* KO mice increased phosphorylation of AMPK^Thr172^ when raised on a HFD, and were resistant to celastrol. Moreover, specific knock-out of AMPK in skeletal muscle largely attenuated the anti-obesity effects of celastrol, supporting the notion that activation of *AMPK* in skeletal muscle is a key step in anti-obesity effects of celastrol.

As a result of increased fatty acid oxidation and decreased glycolysis, FFA levels were reduced dramatically in L6 cells upon celastrol treatment. Importantly, circulatory FFAs were decreased in DIO mice treated with celastrol or in *Pfkm* KO mice. In obese animals, fatty acid oxidation levels were decreased and glycolysis levels were increased in skeletal muscle, which elevated circulating FFAs. Using the 293t/hLepRb model, treating cells with palmitic acid significantly attenuated cellular leptin response. Moreover, boosting cellular fatty acid oxidation by AICAR treatment or glucose starvation abolished this PA-induced leptin desensitization, whereas inhibiting CPT-1 by etomoxir abolished the effects of AICAR and glucose starvation on leptin signaling. However, inhibition of CPT1 in hypothalamus successfully suppresses food intake^38^. Since hypothalamus only uses glucose and ketone as energy sources, inhibition of CPT1 might not reduce FFAs, but rather sense circulating FFAs. In summary, our work supports that a functional metabolic state is necessary for leptin signal transduction, and that celastrol promotes systematic sensitization to leptin signaling by reducing circulatory FFAs.

The SOCS family of proteins has long been considered key negative regulators of leptin signal transduction. Besides SOCS3, SOCS1 also binds and inhibits the JAK family^39,40^, and the expression of SOCS1 is increased in skeletal muscle and the hypothalamus of obese animals^41,42^. However, it has been shown that leptin administration induces SOCS3 but not SOCS1 in the hypothalamus^43^. Our work demonstrated that the metabolic state could influence leptin signal transduction by regulating SOCS1 expression in both 293t/hLepRb cells and mouse skeletal muscles. In 293t/hLepRb cells, in correlation with diminishing leptin signaling, PA treatment reduced SOCS1 expression, and knocking down *SOCS1* abolished PA-induced leptin desensitization. Moreover, celastrol treatment largely reduced *SOCS1* mRNA levels in wild-type but not in *AMPK* mKO skeletal muscles. Therefore, induction of SOCS1 expression by circulatory FFAs is involved in obesity-associated systemic leptin resistance. Since both IL1R1 and SOCS1 play important roles in leptin sensitization and inflammation^15^ ^44^, crosstalk between these two processes may be an important issue to be investigated.

PFKM is the only phosphofructokinase in muscle. In contrast, PFKM is expressed in the brain and most other tissues express PFKL and PFKC as well, forming homo- and hetero-tetramers^24^. It has been reported that mice lacking PFKM in the brain but not muscles have greatly reduced fat stores under chew diet ^45^. This work indicated a close relationship between activity of central PFKM and bodyweight regulation. However, the effects of peripheral PFKM on body weight regulation and whether these effects still maintained in obesity models have not been clarified. In *Pfkm* KO mice we generated (complete loss of PFKM), no obvious differences in behavior and development were observed in comparisons to wild-type littermates, except for anti-obesity effects when fed a HFD. Importantly, *Pfkm* KO mice are completely resistant to celastrol. In contrast, specifically disrupting AMPK in muscles attenuated the anti-obesity effects of celastrol, and weakened celastrol-induced leptin sensitization in liver, hypothalamus, and skeletal muscles. Considering that skeletal muscle accounts for 40% of bodyweight and consumes more than 30% of oxygen in the rest state^46^, metabolic switching in skeletal muscle could systematically improve leptin signal transduction in multiple tissues through metabolites like FFAs.

As the most potent anti-obesity agent, celastrol is a promising drug candidate for treating obesity in humans. However, celastrol is a triterpene that contains an electrophilic quinone methide moiety, which enables celastrol to covalently bind to the cysteine of target proteins. Moreover, as a complicated nature product, the pharmaceutical properties of celastrol are difficult to improve by modifying the compound. Thus, despite the dramatic leptin sensitizing effects, the used of celastrol as an anti-obesity drug is severely limited by resulting adverse side effects^30,31^. Here we discovered that celastrol’s anti-obesity function depends on its ability to inhibit PFKM. Based on key anti-obesity effects on targeting PFKM, we identify a lead compound, 3-79, which efficiently inhibited PFKM activity. Importantly 3-79 reduced body weight of DIO mice as efficiently as celastrol with minimal side-effects. All of this indicated inhibition of PFKM by small molecular holds great promise as a future therapeutic intervention to cure obesity.

## Acknowledgments

We thank the Metabolomics Center, Proteomics Facility and Chemistry Center at the National Institute of Biological Sciences, Facility Center of Metabolomics and Lipidomics at Tsinghua University and Beijing Omics biological technology Co., Ltd. for technical assistance. We thank Dr. Li Li and Dr. Dapeng Ju for helpful discussions. We thank Dr. D. O'Keefe for helpful discussions and comments on the manuscript. This work was supported by grants from the National Natural Science Foundation of China (81670891 and 81870693) awarded to T. Wang, and National Major Scientific and Technological Special Project for “Significant New Drug Development” during the Twelfth Five-year Plan Period (2013ZX0950910) awarded to Z. Zhang.

## Supplemental Figure legends

**Figure S1. 1-85 has equal leptin sensitization and weight losing effect to celastrol. Related to Figure 1**.

(A-B) 1-85 had equal weight losing effect to celastrol in wild-type DIO mice. During 20 days treatment, both 1-85 (0.15 mg/kg) (from 43.07±3.60 g to 36.81±2.80 g, n=7) and celastrol (0.15 mg/kg) (from 42.97±2.27 g to 35.84±1.43 g, n=7) reduced body weight of DIO mice significantly, whereas the body weight of vehicle treated mice (from 43.85±3.75 g to 40.73±3.87 g, n=5) showed no statistically significant decrease. (C) 1-85 had same leptin sensitization effect as celastrol. The eight weeks old wild-type lean mice (n=5 for each group) were treated with vehicle, celastrol (0.15 mg/kg) or 1-85 (0.15 mg/kg) for three days, and subsequently received either saline or leptin (5 mg/kg). The body weight was recorded. The body weight changes 24 hrs after leptin administration were quantified. Vehicle+Saline, −0.02±0.51 g; Vehicle+Leptin, −0.24±0.32 g; Celastrol+Saline, −0.48±0.26 g; Celastrol+Leptin, −1.28±0.08 g; 1-85+Saline, 0.40±0.49 g; 1-85+Leptin, −1.16±0.63 g.

**Figure S2. PFKM expression are successfully blocked by shRNA. Related to Figure 1**.

Knockdown efficiency of *Pfkm* shRNA in L6 cells was examined by western blot.

**Figure S3. The schematic diagram of *Pfkm* KO mice. Related to Figure 1**.

(A) The schematic diagram of double-strand DNA donor generation. (B) Editing sites of Crispr-cas9 system in *Pfkm* locus. (C) The critical exon 5/6 is flanked by two loxP sites in *Pfkm* KO mice. (D) Relative *Pfkm* mRNA level of *Pfkm* KO mice and wild-type control in skeletal muscle measured by QPCR.

**Figure S4. The serum leptin levels of DIO mice. Related to Figure 1 and 2**.

(A) The serum leptin levels of *Pfkm* KO DIO mice (21.00±21.87ng/mL, n=4) were significantly lower than wild-type DIO littermate (71.94±13.60ng/mL). (B) The serum leptin levels of *AMPK*α*1*α*2* mKO DIO mice (82.93±5.55ng/mL) were equal to wild-type DIO mice (86.95±10.11ng/mL).

**Figure S5. Loss of PFKM does not activate AMPK in both liver and hypothalamus. Related to Figure 1 and 2**.

(A-B) Western blots show that both *Pfkm* KO and celastrol treatment does not affect phosphorylation of AMPK^Thr172^ and ACC^Ser79^, but induce phosphorylation on STAT3^Tyr705^ in both liver (A) and hypothalamus (B).

**Figure S6. Phosphorylation on STAT3^Tyr705^ is induced by leptin in both central and periphery tissues. Related to Figure 2**.

Wild-type lean mice at age of eight weeks old were subjected to acute intraperitoneal leptin administration (5 mg/kg), and phosphorylation of STAT3^Tyr705^ were measure in both hypothalamus and skeletal muscles by western blot.

**Figure S7. Celastrol reduce glycolytic rates in 293t/hLepRb cells. Related to Figure 3**.

(A) Celastrol inhibits PFKL at high concentration. The IC50 of celastrol was measured on recombinant PFKL. (B) Seahorse glycolysis stress test of 293T/hLepRb cells treated with celastrol.

**Figure S8. SOCS1 are knocked down by shRNA. Related to Figure 3**.

Knockdown efficiency of two *SOCS1* shRNAs in 293T/hLepRb cells was examined by western blot against SOCS1.

**Figure S9. 3-79 does not reduce body weight of regular lean mice. Related to Figure 4**.

Body weight (g) of eight-week old lean mice (n=5 for each groups) treated with vehicle or 3-79 (1 mg/kg) twice a day for 7 days.

**Table S1. Protein list of Celastrol’s target detected by ABPP in mice skeletal muscle. Related to Figure 1**. Only the proteins with more than 1000 Mascot score in probe group but not in control group were picked.

**Table S2. Targeted metabolimics analysis in celastrol treated L6 cells. Related to Figure 2**.

**Table S3. The levels of FFA in L6 cells. Related to Figure 2**. The L6 cells were treated with 0 μM, 2 μM or 4 μM celastrol for 30 min.

**Table S4. FFA levels in serum of DIO mice. Related to Figure 3**. The DIO mice of wild type and *Pfkm* KO were subjected to daily intraperitoneal treatment of vehicle or 0.15 mg/kg celastrol for 20 days.

**Table S5. Structure and IC50 of original PFKM inhibitor candidates identified by High Throughput Screening. Related to Figure 4**.

