## Supporting info for "Celastrol directly inhibits PFKM to induce weight loss and leptin sensitization"

**SUPPLEMENTAL EXPERIMENTAL PROCEDURES**

**Reagents**

Celastrol (CAS: 34157-83-0) was purchased from Micxy Reagent (Chengdu, China). Withaferin A (CAS:5119-48-2) was from J&K Chemicals (Beijing, China). (+)-Etomoxir (CAS:828934-41-4) was from Target Mol (Shanghai, China). All the other chemical reagents were purchased from Heowns (Tianjin, China) and Ouhe (Beijing, China). Protease inhibitor cocktail (Stock: 04693132001), Phosphatase inhibitor cocktail (Stock: 4906845001) were from Roche (Basel, Switzerland). Streptavidin Agarose (Stock: 20359) was from ThermoFisher Scientific (Waltham, USA). Silver staining kit (Stock: PROTSIL1-1KT) was from Sigma-Aldrich (St. Louis, USA). Vigofect (Stock: T001) was from Vigorous Biotechnology (Beijing, China). Ni Resin (Stock: 88221) was from ThermoFisher Scientific (Waltham, USA). Triose phosphate isomerase (Stock: T6258), aldolase (Stock: 8811) and α-glycerophosphate dehydrogenase (Stock: G6751) were from Sigma-Aldrich (St. Louis, USA). Mouse Pfkm (Stock: TRCN0000012536), human SOCS1 (Stock: TRCN0000057065, TRCN0000057066) TRC shRNA were from Sigma-Aldrich (St. Louis, USA). Seahorse glycolysis stress kit (Stock: 103020-100), microplates, sensor cartridges (Stock: 102905-100) and all related reagents were from Agilent Technologies (Santa Clara, USA). HPBCD (Stock: 346111107B) was from Roquette Freres (Lestrem, France). CollagenⅠ,Rat Tail (Stock: 354236) was from Corning (Corning, USA). Mouse Leptin (Stock: cyt-351-c) was from ProSpec (Ness-Ziona, Isreal). Leptin ELISA kit (Stock: 90030) was from Crystal Chem (Elk Grove Village, USA). HSP90 (Stock: 4877T), AMPK (Stock: 2532), AMPKThr172 (Stock: 2535T), ACC (Stock: 3676T), ACCSer790 (Stock: 11818S), STAT3Tyr705 (Stock: 9139T), HRP-linked anti-rabbit IgG (Stock: 7074S) antibodies were from Cell Signaling Technology (Danverse, USA), PFKM (Stock: GTX111597) antibodies was from GeneTex, Inc (Irvine, USA). Enhanced chemiluminisence substrate (Stock: NEL105001EA) was from PerkinElmer (Waltham, USA). West Femto Maximum Sensitivity Substrate (Stock: 34095) was from ThermoFisher Scientific (Waltham, USA). RNAiso Plus (Stock: 9108), TB green qPCR mix (Stock: RR420A) and Guide-it sgRNA In Vitro Transcription Kit (Stock: 632635) was from Takara Bio (Kyoto, Japan). FastPfu DNA Polymerase (Stock: AP221) and cDNA synthesis kit (Stock: AT341-01) was from TransGen Biotech (Beijing, China). Restriction endonucleases, T4 ligase (Stock: M0202) were from NEB (Ipswich, USA). PMSG (Stock: 493-10-2.5) was from Lee Biosolutions (Maryland Heights, USA). Recombinant Cas9 protein (Stock: 1081059) was from IDT (Coralville, USA).

**Cells and cells culture**

L6 cells and 293t/hLepRb cells were obtained from National Infrastructure of Cell Line Resource, China. Both cells were cultured in 5% CO_2_ incubator in 10 cm^2^ cell culture dishes with DMEM, which contained 10% fetal bovine serum, 1% penicillin-streptomycin complex. In all the experiment using 293t/hLepRb cells, the culture plate was pre-coated with CollagenⅠfrom rat tail.

**Animals**

This study was performed in strict accordance with the recommendations in Guide for the Care and Use of Laboratory Animals of the National Institutes of Biological Sciences, Beijing. All of the animals were handled according to the guidelines of the Chinese law regulating the usage of experimental animals and the protocols (M0020) approved by the Committee on the Ethics of Animal Experiments of the National Institute of Biological Sciences, Beijing. Mice were group housed (up to 5 animals per cage) on a 12:12 hour light-dark cycle, with free access to food and water in individually ventilated specific pathogen free (SPF) cages. All mice used were healthy and were not involved in any previous procedures nor drug treatment unless indicated otherwise. *CKM-CreER* mice (Stock: 006475), *AMPKα1^fl/fl^* mice (Stock: 014141), *AMPKα2^fl/fl^* (Stock: 014142) are from Jackson Laboratories.

***Pfkm* knockout mice generation**

The *Pfkm* KO mice were generated by inserting two cis loxP repeats in the intron 5’ to the exon 5 and 3’ to the exon 6. The schematic diagram of *Pfkm* KO mice was presented in Figure S3. Briefly, to generate double-stranded DNA donor containing exon 5, exon 6 and loxP sequence, the 5’ homologous arm, 3’ homologous arm and cassette containing loxP sequence were amplified from mice genome separately. The primers used for 5’ homologous arm amplification were 5’-ACGCgtcgacGGCTTACATTTGCCCTTTACTCTTTGC-3’ and 5’-GATaagcttcgactcgTCATCTAGCTCATTGGACATTTAATCATCA-3’. The primers used for 3’ homologous arm amplification were 5’- ATATactagtcaacgTCCGGGGTCCTGAATACCTGGG-3’ and 5’- GGAAAAgcggccgCCCCACAGGACAGAGAGGTGACAAG-3’. To generate the cassette containing loxP sequence, the DNA fragment was first amplified from mice genome using primer pairs 5’- TGTATGCTATACGAAGTTATgctgctTCTCGGTAGGCCCCTAGGGAGTGT-3’ and 5’- CATACATTATACGAAGTTATggtatccGCAACATGCCTAAGGACAGGCTGG-3’. Then this fragment was used as template and amplified using primers with sequence as 5’- GATaagcttATAACTTCGTATAATGTATGCTATACGAAGTTATgctgctTCTC-3’ and 5’- ATATactagtATAACTTCGTATAGCATACATTATACGAAGTTATggtatccGCAAC-3’ to generate the final fragment containing loxP sequence. These three fragments including 5’ homologous arm, loxP cassette and 3’ homologous arm were integrated into pBluescript II plasmid, and the plasmids were digested with SalI and NotI to drop out the double-stranded DNA donors before injection (Figure S3A).

The Guide-it sgRNA In Vitro Transcription Kit was used to generate two gRNAs. The two forward primers to generate gRNAs were 5’-TTAATACGACTCACTATAggagctagatgatctcggtGTTTAAGAGCTATGCTGGA-3’ and 5’-TTAATACGACTCACTATAggtccttaggcatgttgctccgGTTTAAGAGCTATGCTGGA-3’, while the reverse primer and template were provided by the kit (Figure S3B). After PCR amplification, the gRNA template was directly used as template for *in vitro* transcription to generate gRNA according to the manual. The gRNA was purified using Guide-it IVT RNA Clean-Up Kit, and immediately used for zygotes injection.

8-weeks-old female C57/BL6J mice were treated with 10 IU PMSG (Pregnant Mare Serum Gonadotropin) and subsequently mated with male C57/BL6J mice. The zygotes were collected and washed with M2 medium. The recombinant Cas9 protein, gRNAs and double-stranded DNA donors were mixed with final concentrations of 80 ng/μL, 20 ng/μL and 20 ng/μL, respectively, followed by injection into zygotes. The zygotes were transplanted into pseudopregnancy ICR mice to generate *Pfkm* KO mice.

Positive founder mice with two loxP sites flanking the exon 5/6 of *Pfkm* locus were screened using PCR and DNA sequencing (Figure S3C). The two loxP sites of *Pfkm* KO mice were determined by PCR. The primers of first loxP site were 5’- GAGGAAGCACAACTACCAAGCACACTG-3’ and 5’- CAATGCCACTCATGTAAGTTCCCTGAC-3’, and the primers of second loxP site were 5’- GGAGGCAGTAGGAAAGTAGAAGCAAGAAC-3’ and 5’- AAGGGTCGCAGTGTCAGGAAGGC-3’. The full fragment between two loxP sites was amplified using primers as 5’- GAGGAAGCACAACTACCAAGCACACTG-3’ and 5’- AAGGGTCGCAGTGTCAGGAAGGC-3’, the PCR product was sent to Sangon Biotech (Beijing) for sequencing.

The mice carrying heterozygous loxP insertion were mated with wild-type C57BL/6J mice and then inbreeded for two generations to avoid mutations through off-target of Crisp/cas9. Both mRNA and proteins of PFKM were abolished in multiple tissues of homozygous *Pfkm* loxP insertion mice including muscle, liver and hypothalamus but not their heterozygous and wild-type littermate (Figure 1J and S3D). To investigate whether this *Pfkm* knockout was caused by loxP insertion or the potential off-target of Crispr-cas9 editing, we performed whole genome sequencing on mice carrying biallelic insertion of loxP (*Pfkm* KO) and their homozygous wild-type littermates. No mutations were confirmed in a 2 Mbp region of *Pfkm* locus. Therefore, Loss of PFKM in *Pfkm* KO animals is caused by loxP insertion rather than the off-target of Crispr-cas9 editing.

**Whole Genome Sequencing**

The whole genome sequencing was performed on one mouse carrying biallelic insertion of loxP in *Pfkm* and its homozygous wild-type littermate. Genomic DNA (4 to 5 μg) was sheared to 300 to 400bp using a Covaris S220 (Covaris, Woburn, MA, USA) and  purified with 1X magnetic beads (Ampure XP; Beckman Coulter). Sheared DNA was subjected to Illumina paired-end DNA library preparation and PCR- amplified for three cycles, and libraries were size selected with 0.55-1X magnetic beads (Ampure XP; Beckman Coulter). Amplified libraries were sequenced using the HiSeq X ten platform (Illumina) as paired-end 150 base reads according to the manufacturer’s protocol.

The sequencing raw reads were aligned to GRCm38/mm10 genome by using BWA-MEM (v0.7.17-r1188). Alignment was done with default parameter settings. Samtools (v1.9) was used to filter mapped reads with the following parameters: -F 1804 -f 2, and the mapped reads were sorted based on genome position.

**Chemical synthesis**

To synthesize 1-85, celastrol (56 mg, 0.125 mmol), EDCI (29 mg, 0.15 mmol), DIEA (35.5 mg, 0.275 mmol) and HOBT (25.25 mg, 0.1875 mmol) were dissolved in 10 mL dried DMF under nitrogen. The propargylamine (8.25 mg, 0.15 mmol) was added dropwise under mild stirring. The solution was kept at room temperature overnight, and then 20 mL ethyl acetate was added into it. This solution was washed three times with water and dried over anhydrous Na_2_SO_4_. After concentration, the resulting residue was purified by silica gel column chromatography with petroleum ether/ethyl acetate (2:1) as the eluent to afford the product.

To synthesize 3-79, 2-acetylthiophene (12.6 g, 0.1 mol) was dissolved in 10 mL acetic acid and heated to 90 ℃. Thirteen mL of 69% nitric acid was mixed with 10 mL acetic acid, followed by added in one portion under stirring. A small amount of sodium nitrate was added, and the exothermic reaction finished in several minutes. Two-hundred mL of water was added to cause separation of a yellow solid, and the solid was washed with aqueous sodium carbonate and recrystallized by methanol.

**Activity-Based Protein Profiling**

Eight-week-old wild-type mice were executed by cervical dislocation, and the soleus, liver, small intestine, stomach, hypothalamus and adipose tissue were removed and homogenized immediately in 10 volume of lysis buffer (50 mM Tris-HCl, pH 7.4, 150 mM NaCl, 1% Triton X-100, 0.5% deoxysodium cholate, 1 mM EDTA, 0.2 mM Na_3_VO_4_, 1 mM PMSF, 1× Protease inhibitor cocktail, 1× Phosphate inhibitor cocktail) by FastPrep-24 homogenizer (MP Biomedicals, Irvine, USA) for 3 times with 40 s on/5 min off cycle at 5 g. The homogenates were centrifuged at 15,000 g, 4 ℃ for 20 min, and the supernatants were collected and the protein concentration was adjusted to 2 mg/mL. The supernatants were divided equally as control and probe groups. For control group, the supernatants were incubated with 200 μM celastrol for 10 min, and were subsequently incubated with 20 μM 1-85 for 60 min at room temperature. For the probe group, the supernatants were incubated directly with 20 μM 1-85 for 60 min. Five volumes of methanol were added to the reaction, and the mixture was kept in -80 ℃ for 60 min for protein precipitation. The mixtures were centrifuged at 2,000 g for 5 min to get sediment. The sediments were washed with methanol for three times, and redissolved in 1 ml 1% SDS aqueous solution. One mM CuSO_4_, 1 mM sodium ascorbate, 0.1 mM TBTA and 0.1 mM 2-azidoethan-1-amine conjugated biotin were added, and the solutions incubated for 120 min at room temperature. Five volumes of methanol were added and the mixture was kept in -80 ℃ for 60 min for protein precipitation. The mixture was centrifuged at 2,000 g for 5 min to get sediment, and the sediment was redissolved in 200 μL 2% SDS aqueous solution by ultrasonic treatment and diluted to 4 ml with deionized water. One-hundred μL streptavidin agarose suspension was added and incubated at room temperature for 120 min, and agarose beads were collected by centrifugation at 2,000 g at 4 ℃ for 5 min and washed with wash buffer (437 mM NaCl, 2.7 mM KCl, 10 mM Na_2_HPO_4_, 2 mM KH_2_PO_4_, 1% NP-40) six times. The agarose beads was boiled with 2×protein loading buffer for 10 min, and 10 μL of supernatant was loaded to 4%-12% SDS-PAGE gradient gel. The electrophoresis was performed at 140 V for 90 min, and the protein bands were detected by silver staining.

**Tandem Mass Spectrometry (MS/MS)**

Protein bands on the SDS-PAGE gels were de-stained and in-gel digested with sequencing grade trypsin (10 ng/μL trypsin, 50 mM ammonium bicarbonate, pH=8.0) overnight at 37 °C. Peptides were extracted with 5% formic acid/50% acetonitrile and 0.1% formic acid/75% acetonitrile sequentially. The peptides extracted were separated by an analytical capillary column (50 μm × 10 cm) packed with 5 μm spherical C18 reversed phase material (YMC, Kyoyo, Japan). Waters nanoAcquity UPLC system (Waters Corp, Milford, USA) was used to generate the following HPLC gradient: 0-30% B in 40 min, 30-70% B in 15 min (A = 0.1% formic acid in water, B = 0.1% formic acid in acetonitrile). The eluted peptides were sprayed into LTQ ORBITRAP Velos mass spectrometer (ThermoFisher Scientific, Waltham, USA) equipped with a nano-ESI ion source. The mass spectrometer was operated in data-dependent mode with one MS scan followed by four CID (Collision Induced Dissociation) and four HCD (High-energy Collisional Dissociation) MS/MS scans for each cycle. Database searches were performed on an in-house Mascot server (Matrix Science Ltd., London, UK) against IPI (International Protein Index) mouse protein database. The search parameters are 10 ppm mass tolerance for precursor ions; 0.7 Da mass tolerance for product ions; two missed cleavage sites were allowed for trypsin digestion. Methionine oxidation was set as variable modification. The search results were filtered with both peptide significance threshold and expectation value to be below 0.05.

**PFK purification and enzymatic inhibition assay**

The his-tag PFK was transiently transfected into 293t cells using VigoFect. 48 h after transfection, 293t cells were collected and suspended in binding buffer (50 mM Tris-HCl, pH=7.4, 100 mM NaCl, 5 mM EDTA, 0.2 mM Na_3_VO_4_, 1 mM PMSF, 1× Protease inhibitor cocktail, 1× Phosphate inhibitor cocktail). The cells lysates were prepared by sonication for 3×15 s on, 30 s off cycle at 30% of the maximum power using ultrasonic liquid processor (Sonics & Materials, Newtown, USA), and were centrifuged at 15,000 g for 20 min at 4 ℃. The supernatants were collected and the protein concentration was adjusted to 2 mg/mL. Ni resin was added to the supernatant and incubated at 4 ℃ overnight. The Ni resin was collected and washed with wash buffer (20 mM Tris-HCl, pH=8.0, 500 mM NaCl, 20 mM imidazole). The recombinant PFK was eluted by elute buffer (20 mM Tris-HCl, pH=8.0, 500 mM NaCl, 200 mM imidazole), and concentrated by using Amicon Ultra Centrifugal Filter (30-kD molecular weight cutoff; Millipore, Burlington, USA) in stock buffer (50 mM Tris-HCl, pH=7.5, 100 mM KCl, 5 mM MgCl_2_, 5% glycerol).

For the PFK inhibition assay, different concentration of compounds was incubated with PFK in binding buffer (50 mM Tris-HCl, pH=7.5, 100 mM KCl, 5 mM MgCl_2_, 5 mM Na_2_HPO_4_, 1 mM NH_4_Cl) for 10 min at room temperature, and the isometric reaction buffer (50 mM Tris-HCl, pH=7.5, 100 mM KCl, 5 mM MgCl_2_, 5 mM Na_2_HPO_4_, 1 mM NH_4_Cl, 2 mM ATP, 0.4 mM NADH, 0.2 mM AMP, 10 mM fructose-6-phosphate, 10 U triose phosphate, 2 U aldolase, 2 U α-glycerophosphate dehydrogenase) was added, the absorbance at 340 nm was measured every 2 min for 30 min.

**Lentivirus Production**

The sequence of *Pfkm* shRNA was 5’-CCGGGCTATGGATGAGAAGAGATTTCTCGAGAAATCTCTTCTCATCCATAGCTTTTT-3’. The TRC lentivirus based *Pfkm* shRNA (TRCN0000012536, Sigma-Aldrich, St. Louis, USA) plasmid was transiently transfected into 293T cells using VigoFect with pSPAX2 and pMD2.g plasmids (the ratio of shRNA, pSPAX2 and pMD2.g was 5:3:2.). Sixty h after transfection, the lentivirus was harvested, and filtered through 0.22 μM filter. The filter liquor was directly used to infect L6 cells. The *Pfkm* shRNA knockdown efficiency was verified by western blot in L6 cells (Figure S2).

**Seahorse glycolysis stress test**

L6 cells were seeded in 96-well seahorse cells culture plate with cells density of 5,000 cells per well. After 8 h, the L6 cells in *Pfkm* RNAi group were infected with *Pfkm* shRNA lentivirus in the presence of 10 μg/ml of polybrene, while the other groups were infected with empty lentivirus. Thirty-six h after infection, the mediums of all the group were replaced with glucose free seahorse assay medium for 1 h. Celastrol was added, and the ECAR was measured immediately. Thirty min after celastrol addition, the glucose, FCCP, rotenone/antimycin A were added according to the standard seahorse glycose stress test protocol, and the ECARs under different conditions were measured using Seahorse XF Analyzer (Agilent Technologies, Santa Clara, USA).

**DIO mice establishment and compounds treatment**

To generate DIO mice, wild-type, *Pfkm* KO, and *AMPKα1α2* mKO mice were fed 45 kcal% fat diet for 20 weeks right after weaning. The body weight was recorded every week to obtain the weight gain curve. For administering compounds to DIO mice, all compounds were first dissolved in DMSO and attenuated with nine volumes of 30% HPBCD solution to formulate the injection, and leptin was dissolved in saline. Celastrol (0.15 mg/kg) and 1-85 (0.15 mg/kg) were administered once a day by intraperitoneal injection in a constant volume of 5 mL/kg body weight 1 h before the dark cycle. Body weight was recorded every day at the same time with compound treatment. The 3-79 (0.5 mg/kg) was administered twice a day by intraperitoneal injection in a constant volume of 5 mL/kg body weight.

**Leptin sensitization assay**

Eight-week-old lean wild-type and *Pfkm* KO mice were intraperitoneal injected with either vehicle, celastrol (0.15 mg/kg), or 1-85 (0.15 mg/kg) in a constant volume of 5 mL/kg body weight 2 h before the dark cycle for four days. On the third and fourth days mice were treated with either saline or leptin (5 mg/kg) in a constant volume of 5 mL/kg body weight 1 h before the dark cycle. Body weight changes between day 3 and day 5 were calculated.

**Protein expression and phosphorylation analysis in mice tissues**

Mice with various genotypes were executed by cervical dislocation, and the soleus, liver, and hypothalamus were removed and frozen immediately. The protein expression and phosphorylation analysis were performed within three days after dissection. The unfrozen tissues were moved to grinding tubes (Sarstedt Inc, Nümbrecht, German), and ten volumes of lysis buffer (50 mM Tris-HCl, pH 7.4, 150 mM NaCl, 1% Triton X-100, 0.5% deoxysodium cholate, 1 mM EDTA, 0.2 mM Na_3_VO_4_, 1 mM PMSF, 1× Protease inhibitor cocktail, 1× Phosphate inhibitor cocktail) were added. The mixture was homogenized by FastPrep-24 homogenizer (MP Biomedicals, Irvine, USA) for 3 times with 40 sec on/5 min off cycle at 5 g. The homogenates were centrifuged at 15,000 g, at 4 ℃ for 20 min, and the supernatants were collected and the protein concentration was adjusted to 4 mg/mL. Forty μg of protein was loaded to 4%-12% SDS-PAGE gradient gel and electrophoresis was performed at 140 V for 2 h at 4 ℃, followed by transfer to PVDF membrane (0.45 μM pore size, Merck Millipore, Burlington, USA) at 100V for 80 min at 4 ℃. The blot was incubated with 5% milk for 1 h at room temperature and incubated with diluted primary antibody (1:1000) with 5% BSA at 4 ℃ overnight. The membranes were washed 3 times with PBST and incubated with diluted anti-rabbit IgG peroxidase conjugate (1:1000, Cell Signaling Technology) at room temperature for 1 h. After washing 3 times with PBST, the membranes were exposed to enhanced chemiluminisence substrate or the West Femto Maximum Sensitivity Substrate for developing the films.

**Mice serum leptin level measurement**

Mouse blood was drawn and placed at room temperature for 1 h to clot. The mixture was centrifuged for 20 minutes at 2,000 g. The supernatants were collected and kept at -80 ℃ until use. Serums were unfrozen on ice before test, and leptin levels were measured using ELISA (enzyme linked immunosorbent assay).

**Protein expression and phosphorylation analysis of L6 or 293t/hLepRb cells**

To investigate AMPK-ACC activation, L6 cells were seeded in 6-well cell culture plates with a density of 2×10^6^ cells per well the day before treatment with different concentration of celastrol or 3-79 for 30 min. Cells were lysed by 200 μL 1×protein loading buffer and boiled at 98 ℃ for 10 min. The mixtures were centrifuged at 15000 g for 15 min, and 10 μL supernatant was loaded to 4%-12% SDS-PAGE gradient gel. The blots were blocked with 5% milk for 1 h at room temperature, and incubated with 1/1000 5% BSA diluted primary antibody at 4 ℃ overnight. Then the membrane was washed 3 times for 5 min each time with PBST and incubated with diluted anti-rabbit IgG peroxidase conjugate at room temperature for 1 h. Then the membranes were washed 3 times for 5 min each time with PBST and exposed to enhanced chemiluminisence substrate for developing the films.

To investigate the relationship between PA and leptin signal transduction, PA was conjugated to BSA by dissolving 50 mM PA in ethyl alcohol and adding nine volumes of 5% BSA under drastic vortex. Then the pH was adjusted to 7.4 using 1 M sodium hydroxide solution. The BSA control was generated in the same procedure without adding PA. The 293t/hLepRb cells were seeded in 6-well cell culture plate with a density of 2×10^6^ cells per well, and were treated with either DMSO, 2 μM celastrol, 1 mM AICAR or glucose starvation for 8 h, followed by BSA or 500 μM PA treatment and subsequent 1 h treatment of vehicle or 10ng/ml leptin. To investigate if SOCS1 was the mediator in PA induced leptin resistance, 293t/hLepRb cells were seeded in 6-well cell culture plate with a density of 2×10^6^ cells per well, and were transfected respectively with two *SOCS1* shRNA plasmids (TRCN0000057065, 5’-CCGGCTTCCGCACATTCCGTTCGCACTCGAGTGCGAACGGAATGTGCGGAAGTTTTTG-3’; TRCN0000057066, 5’-CCGGGACACGCACTTCCGCACATTCCTCGAGGAATGTGCGGAAGTGCGTGTCTTTTTG-3’.) using VegoFect. After 24 h, the cells were treated with BSA or 500 μM PA for 8 h followed by leptin treatment for 1 h. Cells were lysed by 200 μL 1×protein loading buffer and boiled at 98 ℃ for 10 min. The mixtures were centrifuged at 15,000 g for 15 min, 10 μL supernatants were subjected to western blot analysis.

**qPCR analysis**

For total RNA extraction, cells were washed with PBS and homogenized in 1 mL Trizol reagent. For the total RNA extraction from mice tissues, the tissues were homogenized in 10 volumes of Trizol reagent by FastPrep-24 homogenizer. The mixture was incubated at 25 ℃ for 5 min, 300 μL chloroform was added and shaken vigorously for 15 s. The mixture was incubated at room temperature for 3 min and centrifuged at 12,000 g for 15 min in 4 ℃. The aqueous phase was transferred to a new tube. RNA was precipitated with 300 mL isopropanol for 10 min, and centrifuged at 12,000 g for 15 min in 4 ℃. The pellet was washed with 75 % ethanol twice and resuspended in 20 μL DEPC water.

One mg total RNA was reverse transcribed using a cDNA synthesis kit. The genes expression levels were quantified by real-time PCR using TB green qPCR mix. In 293t/hLepRb, the qPCR quantification of transcripts were normalized to β-actin. In mice muscle, the qPCR quantification of transcripts were normalized to *Rn18s*. The following primers were used: human *β-actin*: 5’-CATGTACGTTGCTATCCAGGC-3’ and 5’-CTCCTTAATGTCACGCACGAT-3’; human *SOCS1*; 5’- CACGCAGCATTAACTGGGATGC-3’ and 5’-TACCCACATGGTTCCAGGCAAG-3’; mouse *Rn18s*: 5’- GCAATTATTCCCCATGAACG-3’ and 5’-GGCCTCACTAAACCATCCAA-3’; mouse *SOCS1*: 5’- GTGGTTGTGGAGGGTGAGAT-3’ and 5’- CCCAGACACAAGCTGCTACA-3’.

**Metabolomics analysis**

After compound treatment, cells were collected and washed with PBS 3 times. The plates was put on dry ice and 2 mL of 80% methanol (pre-chilled to -80 ℃) was added. The mixtures were incubated at -80 ℃ for 2 h, then the cell lysate/methanol mixtures were transferred to a 15 mL tube on dry ice. The plates were washed with 1 mL 80% methanol, and the washing liquor was transferred to the same 15 mL tube. The mixtures were centrifuged at 14,000 g for 20 min at 4 ℃ and the metabolite-containing supernatants were transferred to a new 1.5 mL tube on dry ice. The supernatant was dried to pellet using speedvac (ThermoFisher Scientific Waltham, USA). The pellet was stored at -80 ℃ and redissolved with 100 μL 50% methanol before analysis.

Targeted metabolomic experiments were performed by TSQ Quantiva (ThermoFisher Scientific Waltham, USA). C18 based reverse phase chromatography was utilized with 10 mM tributylamine, 15 mM acetate in water and 100% methanol as mobile phase A and B, respectively. This analysis focused on metabolites of the TCA cycle, glycolysis pathway, pentose phosphate pathway, amino acids and purine metabolism pathway. In this experiment, we used a 25-minute gradient from 5% to 90% mobile B. Positive-negative ion switching mode was performed for data acquisition. Cycle time was set as 1 second and a total of 138 ion pairs were included. The resolution for Q1 and Q3 are both 0.7 FWHM. The source voltage was 3500 V for positive and 2500 V for negative ion mode. The source parameters are as follows. Capillary temperature, 320 ℃; heater temperature, 300 ℃; sheath gas flow rate, 35 SLM; auxiliary gas flow rate, 10 SLM. In the targeted method, the ion transitions of over 300 metabolites were optimized using chemical standards. Tracefinder 3.2 (ThermoFisher Scientific Waltham, USA) was applied for metabolite identification and peak integration.

**Lipidomics analysis**

Mouse serum was incubated at 4 ℃ for 1 h, and 50 μL serum was mixed with 150 μL 50% isopropanol/50% acetonitrile and 20 μg/mL FFA 19:0, and vortexed for 5 min. The mixture was centrifuged at 4 ℃ for 5 min, the supernatant was concentrated 10 times with 50% isopropanol/50% acetonitrile. L6 Cells was harvest and freeze-dried, 1000 μL methanol containing 20 μg/mL FFA 19：0 was added. The mixture was vortexed for 5 min and ultrasonically treated for 2 h. The mixture was centrifuged at 4 ℃ for 10 min, and the supernatant was collected.

The FFAs in collected supernatant were separated by ACQUITYUPLC BEH C8 1.7um 2.1*100 mm chromatographic column (Waters, Milford, USA). A Waters UPLC I-Class system (Waters, Milford, USA) was used to generate the following UPLC gradient. 90% A 0-3 min, 65% A 3-6 min, 0% A 7-11 min, 90% A 11-13 min (A, 60% acetonitrile in water, B, 50% isopropanol/50% acetonitrile). The flow rate was 0.26 mL/min, and the column temperature was at 55 ℃. The eluted FFAs were sprayed into a XEVO TQ-XS benchtop tandem quadrupole mass spectrometer (Waters, Milford, USA) via an electrospray ion source in negative ion model. The mass spectrometer was operated with a capillary voltage of 3.0 kV in negative mode. The capillary temperature was set at 300 °C. Sheath gas flow rate and aux gas flow rate were set at 45 and 10 (in arbitrary units). Aux gas heater temperature was 350 ℃. The Slens rf level was 50.0. The resolutions of 120 000 and 30 000 were set for full scan MS and data dependent MS/MS (ddMS^2^) in both modes. The AGC target and maximum IT were 3x10^6^ ions capacity and 100 ms in full scan MS settings while their values were 1x10^5^ ions capacity and 50 ms in ddMS^2^ settings. The TopN (N, the number of top most abundant ions for fragmentation) was set to 15. The normalized collision energy (NCE) was set 25, 35, 45 eV, respectively. The scan range was set at m/z 133.4-2000. FFA quantitation was achieved by comparing to the interior label of FFA 19:0.

**High throughput screening**

The PFKM enzyme inhibition based high throughput screening was performed on a 6,000 Selective Pilot Screening Library. The 384 well plate was seeded with recombinant hPFKM 10 μg per well in 20 μL binding buffer (50 mM Tris-HCl, pH=7.4, 100 mM NaCl, 5 mM EDTA, 0.2 mM Na_3_VO_4_, 1 mM PMSF, 1× Protease inhibitor cocktail, 1× Phosphate inhibitor cocktail) using Combi Reagent Dispenser (WellMate, ThermoFisher Scientific, Waltham, USA). Different compounds was added by automated HTS facilities (Biomek FXP, Beckman, Brea, USA), two final concentrations of library compound including 1 μM and 10 μM was selected. The mixture was incubated at room temperature for 1 h, followed by addition of 20 μL reaction buffer (50 mM Tris-HCl, pH=7.5, 100 mM KCl, 5 mM MgCl_2_, 5 mM Na_2_HPO_4_, 1 mM NH_4_Cl, 2 mM ATP, 0.4 mM NADH, 0.2 mM AMP, 10 mM fructose-6-phosphate, 10 U triose phosphate, 2 U aldolase, 2 U α-glycerophosphate dehydrogenase). The absorbance at 340 nm was recorded every 10 min using EnSpire Multimode Plate Reader (PerkinElmer, Waltham, USA). The inhibition effect of 10 μM celastrol was defined as 100%, while the DMSO was defined as 0%. The compounds which inhibit PFKM activity more than 50% at both 1 μΜ and 10 μM were selected as inhibitor candidates. The IC_50_ of these candidates was measured in buffer with either 0.5 mg/mL bull serum albumin or 1 mM glutathione to exclude the nonspecific protein binding and oxidization, respectively.

**Statistics**

Statistical analysis and data plotting were performed using Excel, and statistical significance was determined using the Student's t-test. Group data in graphs are shown as the mean ± standard deviation (S.D.). Statistical details for each experiment, including the sample numbers (n), can be found in the corresponding figure legend. P < 0.1 was considered statistically significant. *P < 0.1, **P < 0.05, ***P < 0.01.
