## Supplementary figures and images for "Celastrol directly inhibits PFKM to induce weight loss and leptin sensitization"

### supplemental figure S1

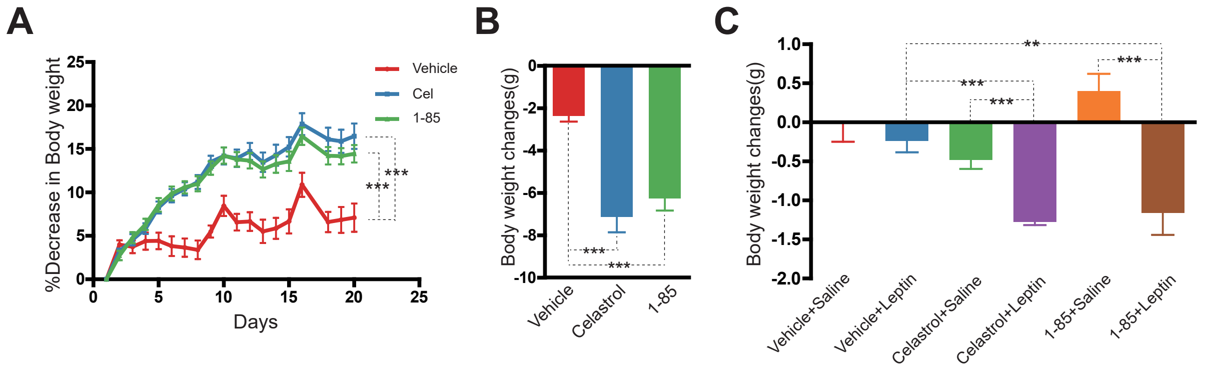

### supplemental figure S2

#### Figure S2

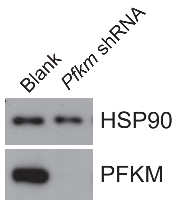

### supplemental figure S3

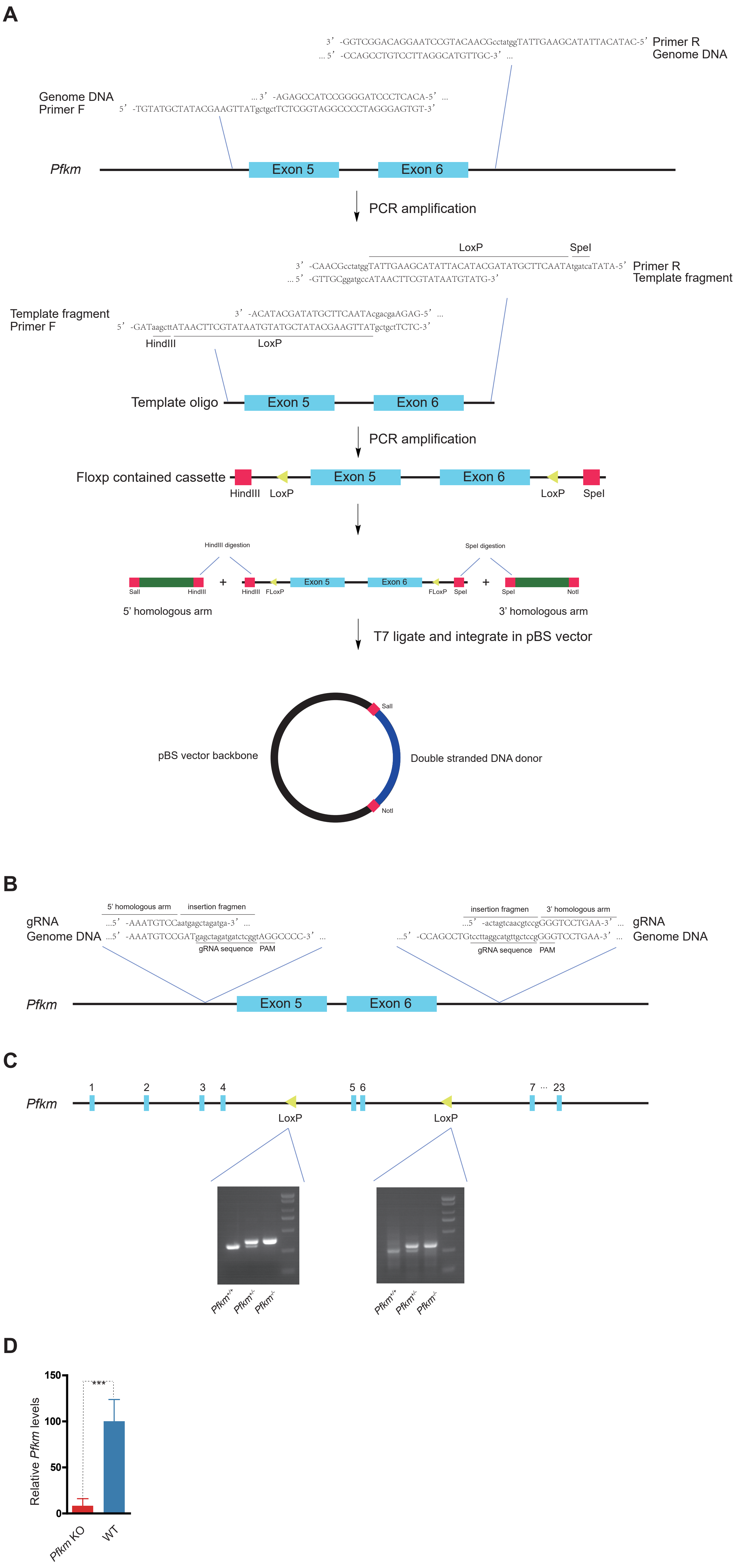

### supplemental figure S4

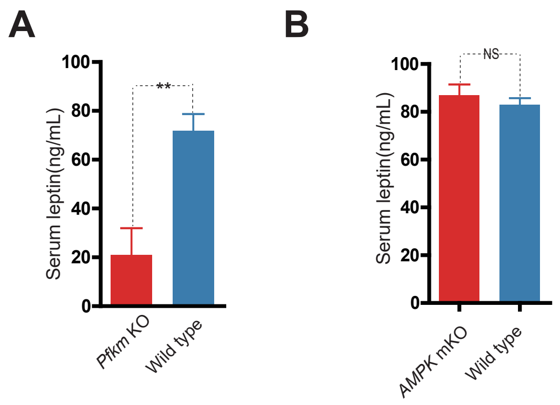

### supplemental figure S5

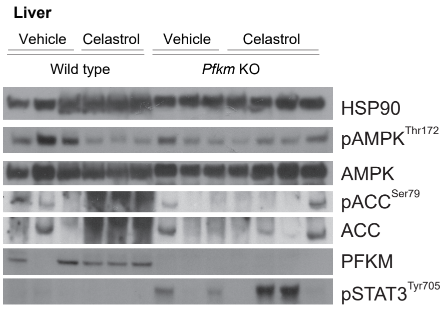

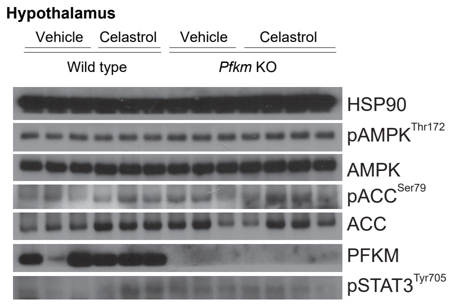

### supplemental figure S6

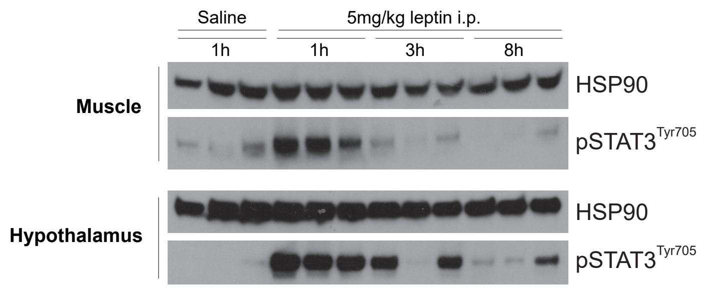

### supplemental figure S7

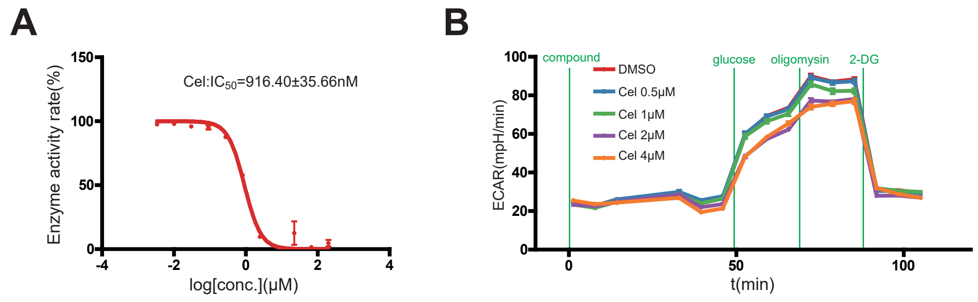

### supplemental figure S8

#### Figure S8

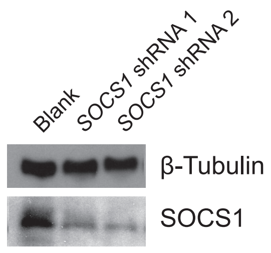

### supplemental figure S9

#### Figure S9

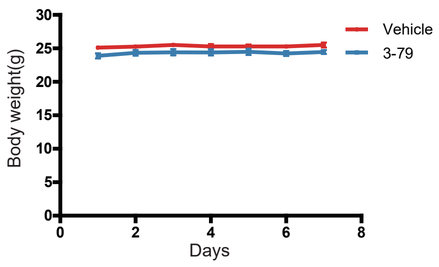
